# An *ex vivo* gut mucosal explant assay to compare HIV tissue susceptibility shows increased HIV susceptibility with methamphetamine exposure

**DOI:** 10.64898/2026.02.05.704074

**Authors:** Julie Elliott, Nicholas E. Webb, Grace D. Cho, Ayub Ali, Peter A. Anton, Otto O. Yang, Jennifer A. Fulcher

## Abstract

**Background:** Existing *ex vivo* mucosal explant models for HIV assess viral replication but do not fully capture the ability of tissue to support productive infection and transfer to target cells, an important indicator of tissue susceptibility. Additional tools to investigate relative mucosal susceptibility to HIV are necessary to better understand risk factors, such as substance use, or to evaluate therapeutics.

**Results:** We developed an *ex vivo* gut mucosal explant assay incorporating a CCR5/CXCR4-expressing GFP-reporter cell co-culture to quantify both viral replication and transfer. Susceptibility was quantified using a composite metric based on time to infection endpoint across viral doses. Modulators of susceptibility, including anti-CD3 stimulation, tenofovir, and methamphetamine, were evaluated. The assay reliably detected differences in tissue susceptibility under various experimental conditions. We detected enhanced susceptibility following anti-CD3 stimulation, while tenofovir pretreatment reduced susceptibility in a dose-dependent manner. Methamphetamine exposure resulted in a modest but significant increase in gut tissue susceptibility to HIV.

**Conclusions:** This novel *ex vivo* explant susceptibility assay advances HIV mucosal transmission research by measuring both viral production and transfer. It can detect subtle changes in susceptibility, providing a versatile platform for evaluating prevention therapeutics, host and microbial factors, and substance use effects.

## Background

Mucosal transmission of HIV is the predominant source of newly acquired infections, and rectal transmission accounts for over 60% of new infections in the United States(1). While most cases of rectal transmission occur in men who have sex with men (MSM), it is an underappreciated source of transmission in heterosexual women(2), making it a phenomenon with broad public health implications.

Methamphetamine use has been repeatedly associated with increased HIV incidence(3-6). Both behavioral and biological factors likely mediate the association between methamphetamine exposure and HIV transmission. Prior studies have shown that methamphetamine can increase expression of HIV co-receptors (CCR5 and CXCR4) and enhance viral replication in CD4+ T cells(7) and myeloid cells(8, 9). At least two clinical studies showed increased inflammatory cytokine expression in the rectal mucosa after methamphetamine use(10, 11); to date no studies have examined the direct impact of methamphetamine on gut tissue HIV infection.

*Ex vivo* gut explant infection models use intact human colorectal tissue with the goal of maintaining cellular architecture and immune microenvironments to study early events of mucosal HIV infection. These models have been used extensively in the pre-clinical testing of HIV prevention therapeutics(12). Colorectal explants also represent an important tool for understanding factors which increase or decrease susceptibility to infection(13); however, standardized methods to compare infection susceptibility across tissues are limited.

Current *ex vivo* explant models use cumulative p24 antigen measured by ELISA or HIV gag measured by quantitative real time PCR as infection endpoints(13-15). These measures quantify viral particle production but do not assess the ability of tissue to produce and transfer productive virus, which is necessary for sustained infection. Further, current models often employ supra-physiologic doses to ensure tissue infection, which limits the ability to compare infectivity across tissues in a physiologic context.

We sought to develop an *ex vivo* explant assay that can quantify relative HIV susceptibility in gut tissues. We approached this by focusing on the ability of gut tissue to support productive viral replication and transfer to susceptible target cells. Here we describe a co-culture system using a limiting dose titration approach with indicator cells to compare relative tissue susceptibility in colorectal tissue explants, and use this method to examine the effects of methamphetamine exposure of gut tissue HIV susceptibility.

## Methods

### Tissue specimens and explant generation

Intestinal tissues used in this study were obtained deidentified from the Translational Pathology Core Laboratory (TPCL), a research facility in the Department of Pathology at the University of California Los Angeles. Normal appearing remnant tissue specimens procured from the large intestine were released for research purposes following pathologist review. Specimens, varying in size from 3cm-10cm^2^, were collected into 25-30ml of RPMI1640 and stored at 4 degrees until transport to the laboratory (generally between 1-12 hours post-surgery). Upon receipt in the laboratory, tissues specimens were washed 3-5 times in sterile PBS and transferred to a petri dish containing explant media (RPMI 1640 supplemented with 10% FBS (Atlanta Biologicals), 1x antibiotic-antimycotic solution (Gibco cat#15240062, Fisher Scientific), and 0.1mg/ml piperacillin-tazobactam (Pfizer, New York, NY). The mucosa was dissected from the sub-mucosa using sterile scissors and 4mm explants were generated using disposable dermal biopsy punches (Integra Miltex cat#12-460-409, Fisher Scientific). Explants were cryopreserved and subsequently thawed according to the protocols detailed by Hughes et al(16). Prior to use, explants were rapidly thawed and transferred using a sterile pipet to a petri-dish containing 15-20ml of explant media for equilibration at room temperature for 10-15 minutes before further use.

### HIV-1 virus expansion

HIV-1_Bal_ was obtained from the NIH AIDS Reagent Program, Division of AIDS, National Institute of Allergy and Infectious Diseases (NIAID). Freshly isolated peripheral mononuclear cells (PBMC) were provided by the UCLA CFAR Centralized Laboratory Support Core then CD4+ T cells were expanded using a CD3-CD8 bispecific antibody in the presence of 50U/ml of IL-2. Following 5-6 days of expansion, cultures comprised >90% CD4+ T cells as verified by flow cytometry. Stocks of HIV-1_BaL_ were generated following incubation for 2-4hrs in 1ml of virus stock and subsequent tissue culture. Virus containing supernatants were harvested between days 6 to 9 and the TCID_50_ determined by Reed-Muench method using activated PBMC. The same viral stock was used throughout this study.

### Indicator cells

T1-CCR5 cells are an X4-HIV-1 permissive immortalized CD4-expressing lymphocyte cell line(17, 18) that was stably transduced with CCR5(19). These cells were then transfected with at Tat- and Rev-dependent eGFP expression gene kindly provided by Dr. Benhur Lee(20). Limiting dilution cloning was performed to enrich for a pure population with high eGFP expression after HIV-1 infection.

### Explant infection and indicator cell co-culture

Explants were thawed and equilibrated as described above. Triplicate explants were added to wells of a 24-well tissue culture plate containing 1ml of complete explant media. Agents proposed to either enhance or reduce mucosal HIV susceptibility were added: anti-human CD3 antibody (12F6 clone, generous gift from Dr. Johnson Wong) 200ng/ml with 50U/ml IL-2 (PeproTech, ThermoFisher cat#200-02-1MG), 10uM or 1uM tenofovir disoproxil fumarate (NIH HIV Reagent Program), 100uM methamphetamine (Sigma cat#M8750), or control (media alone). Cultures were incubated overnight at 37°C /5% CO2 in a humidified tissue culture chamber. The following day, freshly thawed HIV_Bal_ was added to the explant cultures at limited dilutions with final TCID_50_ of 5x10^4^, 5x10^3^, or 5x10^2^ then incubated at 37°C for 2 hours. Pilot experiments included an additional negative control condition with TCID_50_ 5x10^4^ heat-inactivated HIV_Bal_. Explants were then transferred to fresh wells containing 1-2ml of explant media and washed thoroughly 5 times by complete media exchange. Washed explants were then individually added to wells of a 24-well plate containing 1x10^5^ T1-R5-GFP cells in 1ml explant media. Co-culture wells were sampled at daily intervals by the removal of 100ul of media, after ensuring that cells were well resuspended. Cells were transferred to a round bottom tissue culture plate and centrifuged at 400g for 5 minutes in sealed buckets and then fixed by resuspension in 2% paraformaldehyde. Amount of infection (%GFP) was analyzed using a FACSymphony A1 cytometer (Becton Dickenson, San Jose, CA) equipped with a high throughput adaptor. Cells were gated using forward and side scatter and the percentage of GFP expression was quantified as compared to an uninfected T1-R5-GFP cell control. A minimum of 2000 events were analyzed.

### Statistical Analyses

Time to infection endpoint was interpolated from T1-R5-GFP time course data using Python 3.6 (scipy.interpolate). The area under the curve (AUC) from the interpolated time to infection vs TCID50 dose graph was calculated using Python 3.6 (numpy.trapz). Comparisons between treatment conditions and/or groups were analyzed using paired or unpaired t-tests, as appropriate. Data visualization and analyses were performed using GraphPad Prism v10.6.1.

## Results

### Development of tissue susceptibility protocol

To quantify relative tissue susceptibility, we created a measure that would include the ability of infected tissue to produce and transfer virus to adjacent cells rather than simply quantify viral nucleic acids or proteins. In addition, we wanted to include limiting dilutions of virus to better mimic physiologic conditions where differences in tissue infection may be discernable. To do this, we developed a protocol where explants were infected with three TCID_50_ doses and infection was monitored by a highly susceptible GFP-reporter cell line that would amplify early virus, thereby making early stages of explant infection easier to detect (**Figure 1A**). The indicator cell line was T1-CCR5-GFP cells, an immortalized CD4+ lymphoblastoid cell line containing a Tat/Rev-dependent eGFP indicator gene for HIV infection. In addition to the GFP reporter, these cells had endogenous CXCR4 expression and stably transduced CCR5 expression, allowing for use with both X4 and R5 viruses (**Figure 1B**). T1-GFP cells were readily infected with HIV by 7 days post-infection, which was inhibited in the presence of tenofovir (**Figure 1C**).

**Figure 1:**
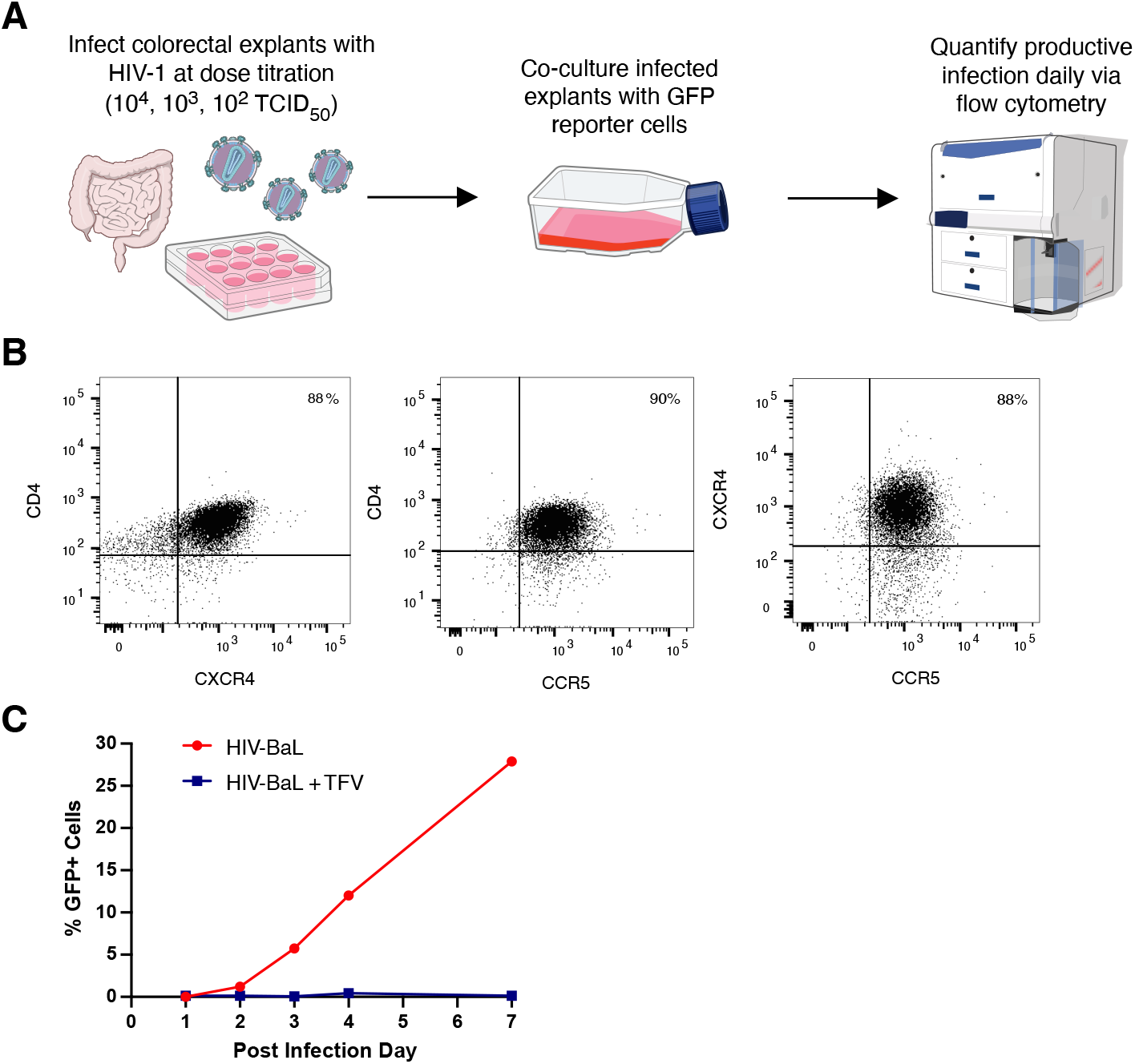
Ex vivo gut tissue HIV susceptibility assay overview. **(A)** Assay schematic using gut explants and GFP-reporter cells to measure productive infection. Illustrations from NIAID NIH BioArt Source. **(B)** CCR5 and CXCR4 co-receptor expression by flow cytometry. **(C)** Infection kinetics of GFP-reporter cell infection using 104 TCID50 HIV-BaL (red). No infection seen when 10uM tenofovir added (blue).

### Infection kinetics differs by input dose and tissue stimulation

The infection kinetics in our co-culture system across doses is shown in **Figure 2A**. As expected, higher TCID50 resulted in early detectable infection with lower TCID_50_ resulting in later detectable infection. Infectability differs by dose so using dose titration composite may give a better assessment of tissue susceptibility. In addition, there is great variability over the course of infection and between individuals. We next increased tissue susceptibility by pre-treating explants with anti-CD3 stimulation with IL-2, then assessed our ability to distinguish differences. As seen in **Figure 2B**, stimulated explants showed an earlier and more rapid increase in measurable T1-CCR5-GFP cell infection. Therefore, our assay can detect differences in infection with or without stimulation, and the greatest differences appear early in the infection.

**Figure 2:**
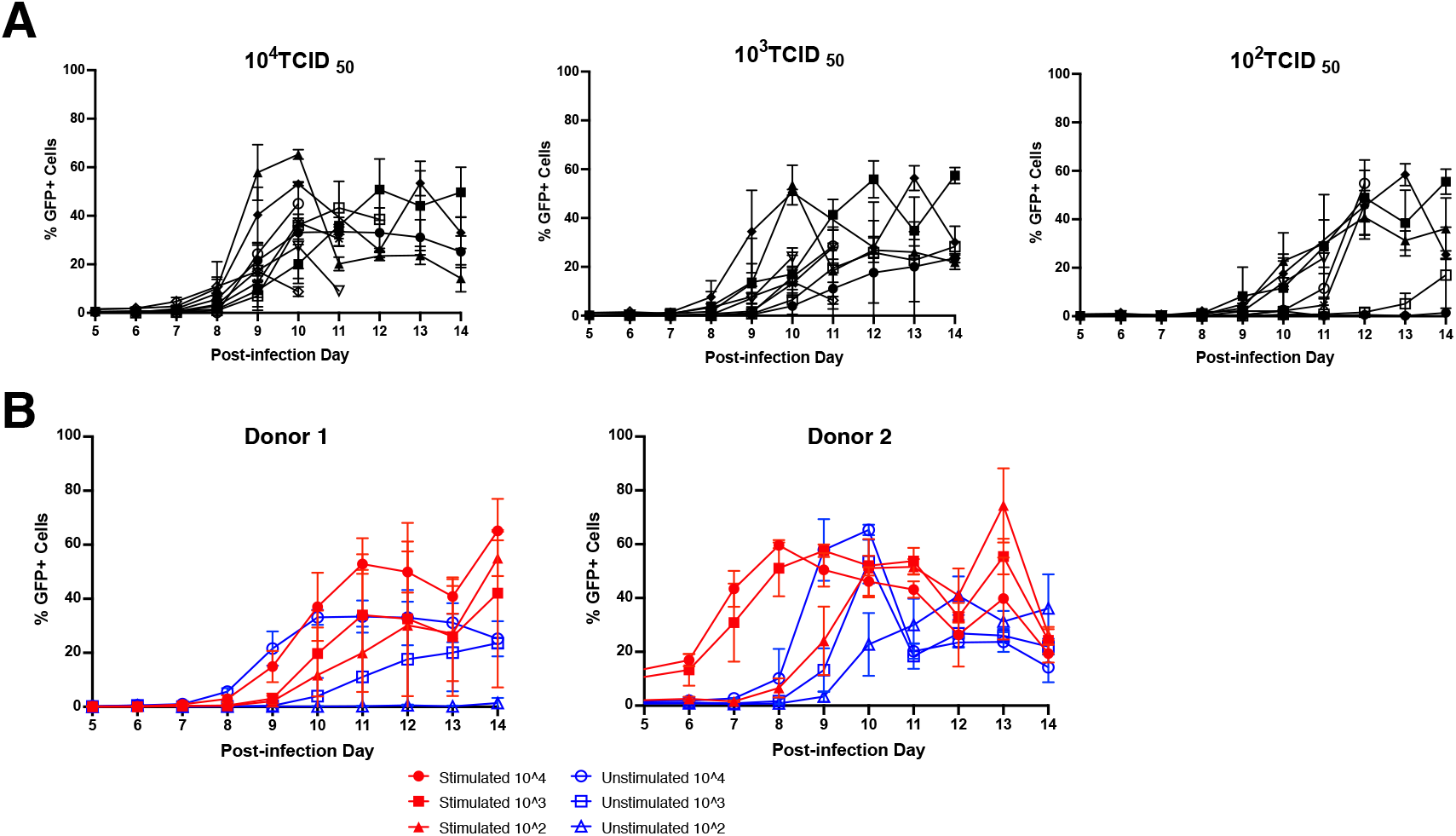
Variation in HIV infectability by virus dose and stimulation. **(A)** Variation of donors by TCID50 dose. Each symbol represents an individual donor; the same 9 donors are included in all three TCID50 doses. Symbols represent the mean % GFP-positive cells from triplicate explant infections with error bars representing standard deviation. **(B)** Effect of anti-CD3 stimulation on infection kinetics.Symbols represent the mean % GFP-positive cells from triplicate explant infections with error bars representing standard deviation.

### Relative susceptibility metric can reliably distinguish changes in HIV susceptibility

In order to compare susceptibility between individuals and/or treatment conditions, we created a metric that incorporated data from all dose titrations. To ensure analysis focused on virus transferred from the explant and not ongoing viral replication in the T1-CCR5-GFP cells, we focused only on the initial phase of infection based on our data (**Figure 2B**). We reasoned that tissues that reached this end point more quickly were more susceptible to infection, and that end point analysis would more reliably reflect the efficiency of the iterative process of infection compared to final amount. Thus, we interpolated the time to infection endpoint defined as 20% GFP-reporter cell infection (**Figure 3A**). When comparing the time to infection endpoint between stimulated and unstimulated tissues, we found inconsistent differences by infection dose (**Figure 3B**). This underscores the importance of using a titration approach to fully assess the tissue susceptibility. In order to incorporate data from all TCID_50_ dose titrations, we used the area under the curve (AUC) for comparisons (**Figure 3C**). As shown in **Figure 4A**, we could reliably detect differences in tissue susceptibility between stimulated and unstimulated conditions in a variety of individuals. We also assessed our ability to detect decreases in susceptibility by pre-treating explants with different doses of the antiretroviral drug tenofovir. As shown in **Figure 4B** we could reliably detect a dose-response decrease in tissue susceptibility with tenofovir using our assay. To make the susceptibility measure more interpretable as a comparative metric, we normalized to the unstimulated condition showing the ability to detect up to 30% differences in susceptibility (**Figures 4C and 4D**). We also compared susceptibility within our tissue donor pool by age and by sex but found no significant differences (**Supplemental Figure 1**) in this small sample group.

**Figure 3:**
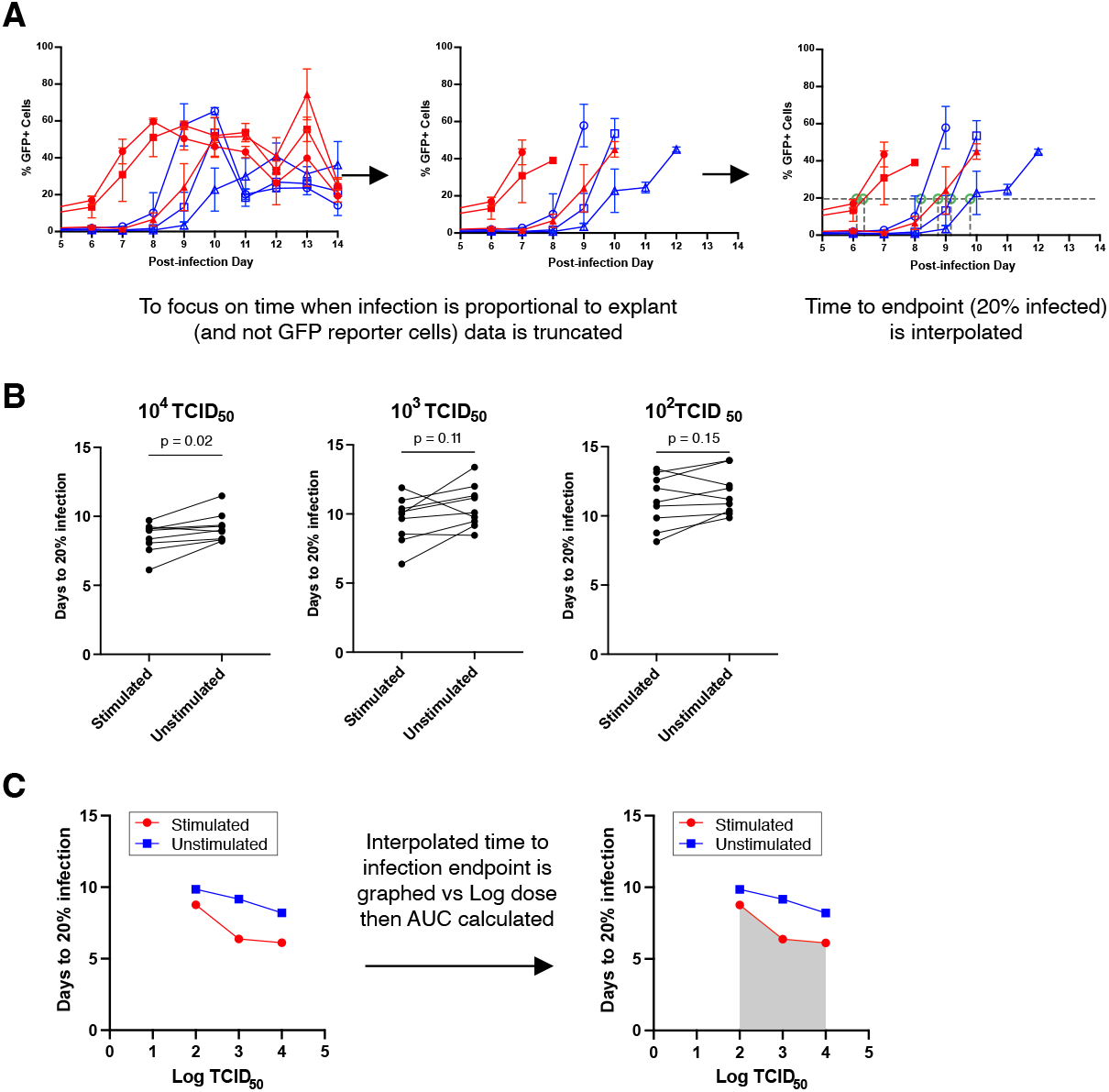
Analytic strategy and composite metric for comparative susceptibility. **(A)**Schematic depiction of infection endpoint interpolation. **(B)** Comparison of interpolated time to infection by TCID50 dose. Each pair of points represents an individual donor. **(C)** Schematic depiction of composite area under the curve susceptibility metric.

**Figure 4:**
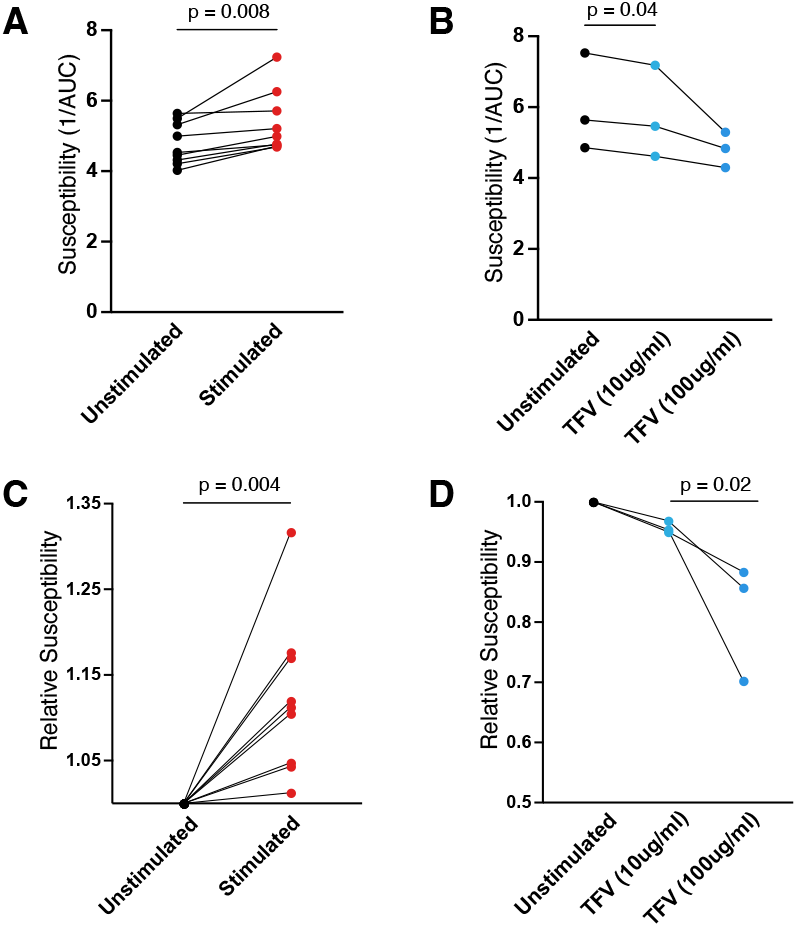
Ex vivo explant susceptibility assay can detect increases and decreases in gut tissue HIV susceptibility. **(A)**Increased susceptibility following anti-CD3 stimulation or **(B)** decreased susceptibility following tenofovir treatment detected by explant susceptibility assay using composite AUC metric. Each paired point represents an individual donor. P-values determined using paired t-tests. Relative susceptibility compared to untreated condition for **(C)** anti-CD3 stimulation and **(D)** tenofovir treatment. Each paired point represents an individual donor. P-values determined using paired t-tests.

### Methamphetamine increases HIV susceptibility in gut tissue

To begin to address a long-standing question regarding the biological effects of methamphetamine on HIV transmission, we examined the direct effects of methamphetamine exposure on HIV susceptibility using our assay. We found that methamphetamine exposure resulted in a small, but significant, increase in HIV susceptibility (**Figure 5**). Coupled with behavioral factors, this measurable change in tissue susceptibility could help explain the increased risk of HIV mucosal transmission in people who use methamphetamines.

**Figure 5:**
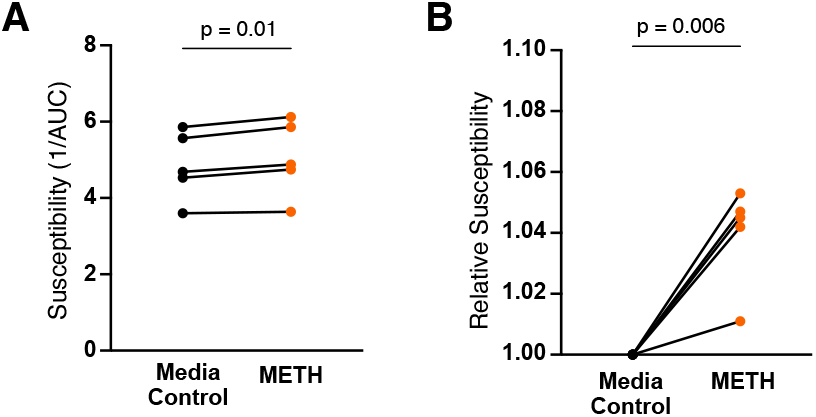
Methamphetamine exposure increases gut tissue HIV susceptibility. **(A)** Increased susceptibility following methamphetamine exposure. **(B)** Relative susceptibility compared to untreated condition following methamphetamine exposure. Each paired point represents an individual donor. P-values determined using paired t-tests.

## Discussion

Here we describe a novel *ex vivo* gut mucosal explant assay to quantify relative tissue susceptibility to HIV infection. This assay goes beyond standard colorectal explant models in that it assesses both virus production and subsequent transfer using a GFP-reporter cell readout. This readout provides a more nuanced assessment of mucosal susceptibility, focusing not only the capacity of tissue to support viral replication but also the efficiency of transmission to target cells. We showed the ability to detect both increases and decreases in relative susceptibility, and used this method to show that methamphetamine can directly increase gut tissue susceptibility to HIV infection.

These findings align with epidemiological data linking methamphetamine use to increased HIV acquisition risk(21, 22), and provides evidence for a biologic mechanism linking substance use and mucosal HIV susceptibility. Previous studies have implicated immune activation as a potential mediators of methamphetamine-associated HIV risk(8, 9). Our study is the first to provide direct *ex vivo* evidence that methamphetamine can act directly on gut mucosal tissue to enhance susceptibility. While these experiments showed potential direct effects of methamphetamine, additional studies are needed to validate these findings and further investigate factors such as duration of the effect, dose-response relationship, and the underlying mechanisms.

A strength of this assay is the ability to detect both subtle and larger increases and decreases in susceptibility. This method could have broad utility investigating host, microbial, and environmental factors that shape mucosal HIV transmission dynamics. The intestinal microbiome is increasingly recognized as a key determinant of mucosal function and inflammation in HIV (23, 24), and more recently has been implicated in HIV susceptibility risk(25, 26). Our assay could be used to evaluate the role of microbiome-derived metabolites or microbial communities in human tissue permissiveness to HIV. In addition, this assay can easily be applied to pre-clinical testing for prevention therapeutics including microbicides or small molecule inhibitors, where quantitative comparisons of relative susceptibility across condition are important.

This method also could be used to examine population-level differences in susceptibility. However, the degree of donor variability inherent in human mucosal explants would necessitate a large sample size to detect significant differences. Nonetheless, the reproducibility of the assay and the ability to detect directional changes make it a potentially valuable comparative tool. Future adaptation for higher-throughput screening using plate-based flow cytometry would increase scalability and enable larger studies of determinants of HIV susceptibility.

This study has several limitations. First, *ex vivo* explant models, while more physiologically relevant than cell line systems, cannot fully recapitulate the dynamic environment of the human gut, including factors such as luminal microbiota, vascular flow, and immune cell trafficking. Second, our assay focuses primarily on productive infection and transmission within a controlled *ex vivo* context, and therefore may not capture the full spectrum of host-pathogen interactions that occur in vivo. Finally, while the assay provides relative susceptibility readouts, it does not directly quantify absolute infection risk.

## Conclusions

Our *ex vivo* gut mucosal explant susceptibility assay enables simultaneous assessment of virus production and transfer, offering a powerful platform to investigate how drugs, microbes, and host factors alter tissue susceptibility. Using this assay we showed that methamphetamine exposure induces a measurable increase in gut mucosal tissue infection; an important finding that helps explain the clinical association between methamphetamine use and increased HIV acquisition. This assay system could be adapted for HIV prevention research, therapeutic development, and to enhance our understanding of the mucosal determinants of HIV acquisition.

## Declarations

### Ethics approval and consent to participate

Tissues used in this study were obtained fully deidentified thus the need for IRB review and/or consent was waived.

### Consent for publication

Not applicable

### Availability of data and materials

The datasets used and/or analyzed during the current study are available from the corresponding author on reasonable request.

### Author Contributions

**JE:** Methodology, Investigation, Writing – Review and Editing; **NEW:** Methodology, Formal analysis, Writing – Review and Editing; **GDC:** Investigation; **AA:** Resources; **PAA:** Conceptualization, Methodology, Writing – Review and Editing; **OOY:** Conceptualization, Methodology, Resources, Writing – Review and Editing; **JAF:** Conceptualization, Methodology, Formal analysis, Visualization, Writing – Original draft, Funding acquisition

## Acknowledgements

Not applicable

